# Specificity and mechanism of 1,6 hexanediol-induced disruption of nuclear transport

**DOI:** 10.1101/2023.03.30.534880

**Authors:** Elizabeth C. Riquelme Barrientos, Tegan A. Otto, Sara N. Mouton, Anton Steen, Liesbeth M. Veenhoff

## Abstract

Selective transport through the nuclear pore complex (NPC) depends on the dynamic binding of the intrinsically disordered components of the NPC, the FG-nups, with each other and with nuclear transport receptors (NTRs). Hydrophobic interactions with the phenylalanines of FG-nups are critical for this dynamic binding. 1,6-hexanediol (1,6HD), is an aliphatic alcohol that interferes with hydrophobic interactions. Here we assessed the specificity and mechanism by which 1,6HD disrupts the permeability barrier of NPCs in live baker’s yeast cells. Exposure to 1,6HD (10 min, 0-5%) leads to gradual loss of the NPC permeability. This is likely a direct effect on the nuclear transport machinery as cell viability, the pH and ATP levels in the cytosol, as well as the appearance of mitochondria, Golgi, peroxisomes, ER, vacuoles, plasma membrane, nucleolus, secretory pathway and stress granules are not notably changed. There are however effects on the cytoskeleton and Hsp104 to be noted. While 1,6HD treatment does not lead to dissociation or degradation of NPC subunits, a massive relocation of multiple NTRs from NPCs does occur. This displacement quantitatively correlates with the increased passive permeability of NPCs. The loss of NTRs and associated cargo will present a major change in the macromolecular crowding and composition and hence the physicochemical properties of the central channel. We conclude that 1,6HD provides a surprisingly specific intervention to temporarily permeate NPCs and we present evidence that the mechanism includes release of NTRs from the NPCs.

## INTRODUCTION

The Nuclear Pore Complex (NPC) is the sole gate between the nucleus and cytosol. The central channel of NPCs is lined with intrinsically disordered phenylalanine-glycine rich nucleoporins, the FG-nups, and it hosts many nuclear transport receptors (NTRs) (Dultz et al. 2022; Hampoelz et al. 2019; Wing, Fung, and Chook 2022; Fernandez-Martinez and Rout 2021). The NTRs bind their cargo and shuttle them through the channel by transiently binding the FG-nups (Paci, Caria, and Lemke 2021; Wing, Fung, and Chook 2022; Bayliss, Littlewood, and Stewart 2000). For the NTR Importinβ it was shown that besides a fraction that is shuttling cargo between the cytoplasm and nucleus, there is also a fraction that is more stably associated with NPCs (Lowe et al. 2015; Kapinos et al. 2014). In addition to NTRs also cargo and non-cargo are present in the NPC. In isolated yeast NPCs, 15,6 MDa worth of NTRs and 10,4 MDa worth of cargo add significantly to the 52,3 MDa mass of actual NPC subunits (Kim et al. 2018). The central channel of the nuclear pore complex is thus a highly crowded and complex environment where the joint presence of NTRs, FG-nups and cargo creates an environment that allows fast and selective transport.

The exact structure of the central channel has remained elusive because experimentally probing its behaviour in living cells is challenging. Our knowledge about the behaviour of the FG-nups and NTRs is inferred from, amongst others, imaging detergent-perforated or live cells (Chowdhury, Sau, and Musser 2022; Schnell, Tingey, and Yang 2022; Mattheyses et al. 2010; Yu et al. 2022), AFM measurements on nuclear envelopes (Sakiyama et al. 2016), transport measurement in biomimetic NPCs (Jovanovic-Talisman et al. 2009; Fisher et al. 2018; Kowalczyk et al. 2011), surface anchored FG-nups (Kapinos et al. 2014) or from probing the structural conformation of purified FG-nups or FG-nup fragment preparations (Frey, Richter, and Görlich 2006; Celetti et al. 2020; Ader et al. 2010; Hayama et al. 2018; Sparks et al. 2018). These experimental studies, together with computational strategies (Davis, Ford, and Hoogenboom 2022; Zheng and Zilman 2023; Isgro and Schulten 2007; Popken et al. 2015; Ghavami et al. 2014), have resulted in a number of models explaining the fast and selective transport through the NPC (Dultz et al. 2022; Hampoelz et al. 2019; Wing, Fung, and Chook 2022; Fernandez-Martinez and Rout 2021; Hoogenboom et al. 2021; Huang and Szleifer 2020). All models agree that the phenylalanines of the FG-repeat regions that are engaging in hydrophobic interactions, as well as the intrinsically disordered nature of the FG-nups, are key parameters. They enable the highly dynamic intra- and inter-chain hydrophobic interactions between FG-repeat regions and with the hydrophobic grooves on the surfaces of NTRs. In the Kap-centric models the slow exchanging pool of NTRs are proposed to be important to create the proper barrier function (Kapinos et al. 2017; Kalita et al. 2022; Fragasso et al. 2022).

Early experiments using aliphatic alcohols pointed to the importance of hydrophobic interactions for import into nuclei of permeabilized cells (Ribbeck and Görlich 2002) and in live yeast cells (Shulga and Goldfarb 2003). Early experiments in permeabilized HeLa cells showed that selective transport of fluorescent reporters (MBP or IBB-MBP) was abrogated in the presence of hexane-1,2-diol but not by the less hydrophobic hexane-1,2,3-triol (Ribbeck and Görlich 2002). In live yeast cells it was observed that the nuclear accumulation of GFP fused to a classical nuclear localisation signal (NLS) was lost upon addition of alcohols and the extend of equilibration was dependent on the hydrophobicity of the alcohol (Shulga and Goldfarb 2003). Biochemical studies using purified FG-repeat fragments show that some of them are cohesive and that their interactions are disrupted by 1,6HD (Patel et al. 2007; Schmidt and Görlich 2015). Also, within the yeast cytosol such overexpressed fragments form foci that are dispersed by 1,6HD (Patel et al. 2007). Lastly, 1,6 HD was shown to increases the diameter of NPCs in *Xenopus* oocyte nuclear envelope preparations (Jäggi et al. 2003). Most dramatically, in the context of mutant NPCs that lack the inner ring nucleoporins Nup170 or Nup188, 1,6HD can even lead to loss of FG-nups from these NPCs (Shulga and Goldfarb 2003; Onischenko et al. 2017). The effect of hexanediol in the above studies was attributed to a reversible disruption of inter-FG repeat cohesion. However, as also the interactions between NTRs and FG-nups are based on hydrophobic interactions, hexanediol will likely also take effect here. Illustrative for the high surface hydrophobicity of NTRs, is their strong binding to a phenyl sepharose chromatography column yielding highly enriched fractions from HeLa cell extracts (Ribbeck and Görlich 2002). Jointly these studies support the importance of hydrophobic interaction for nuclear transport, and the potential of 1,6 HD to disrupt those.

Unrelated to nuclear transport, 1,6HD has also been widely used to dissolve liquid-liquid phase separated compartments in cells and to dissolve condensates in *in vitro* studies. With aggregation-prone peptides, the alcohol dissolves hydrogels (Molliex et al. 2015; Kroschwald, Maharana, and Simon 2017; Shi et al. 2017) but not fibers (Lin et al. 2016; Van Lindt et al. 2022). In cells, the interpretation of effects of 1,6HD are more difficult (Kroschwald, Maharana, and Simon 2017) and depending on the cell type, growth condition and the concentration and length of treatment different results may be obtained. There are many examples of discrepancies in the literature; only one example is the organization of actin and tubulin. While some reports show that they are affected by 1,6HD (Wheeler et al. 2016; Kroschwald, Maharana, and Simon 2017), others report that microtubules are unaffected (Lin et al. 2016).

From the above, the question arises how specific the effects of 1,6HD on nuclear transport are and, whether they are based on a loss of cohesion between the FG-repeat regions, or between FG-nups and NTRs, or both. Here, we probe the impact of 1,6HD on nuclear transport by measuring the effects on passive transport, on NTR-facilitated import and export, and on the cellular localisation of Nups and NTRs. We also assess a large number of possible indirect effects of 1,6HD, namely cell viability, the pH and ATP levels in the cytosol, and the appearance of mitochondria, Golgi, peroxisomes, ER, vacuoles, plasma membrane, nucleolus, secretory pathway, stress granules, the cytoskeleton and Hsp104 foci. Our data support that 1,6HD provides a surprisingly specific intervention to temporarily increase the passive permeability of NPCs by the release of NTRs from the NPC.

## RESULTS

### Disruption of the permeability barrier of NPCs by 1,6 hexanediol

Previous reports already showed that 1,6HD disrupts the permeability barrier of NPCs in yeast cells (Shulga and Goldfarb 2003; Patel et al. 2007). We add to this work and provide a quantitative assessment of the impact of 1,6HD on passive nuclear entry of large reporters and NTR-mediated transport of GFP-NLS and GFP-NES reporters in yeast. To assess passive nuclear entry, the MG5 reporter, composed of a Maltose Binding Protein and 5 GFPs is used. MG5 has a molecular weight of 177 kDa and is excluded from the nucleus in wild type cells (Popken et al. 2015). Mid exponential growing cells were exposed for 10 minutes to zero, 0.625, 1.25, 2.5 or 5% 1,6HD, or to the less hydrophobic alcohol 2,5 hexanediol (2,5HD). The steady state distribution of the GFP-reporters was calculated by taking the ratio of the fluorescence measured in the nucleus and the cytosol (the N/C ratio). The permeability of the NPCs for entry of MG5 increased gradually with increasing concentrations of 1,6HD (Fig 1A) indicating that NPCs became more permeable for this large protein. 1,6HD had a stronger effect on the passive permeability of NPCs than 2,5HD, as MG5 remains properly excluded from the nucleus, even at a concentration of 5% 2,5HD (Fig 1A).

**Figure 1:**
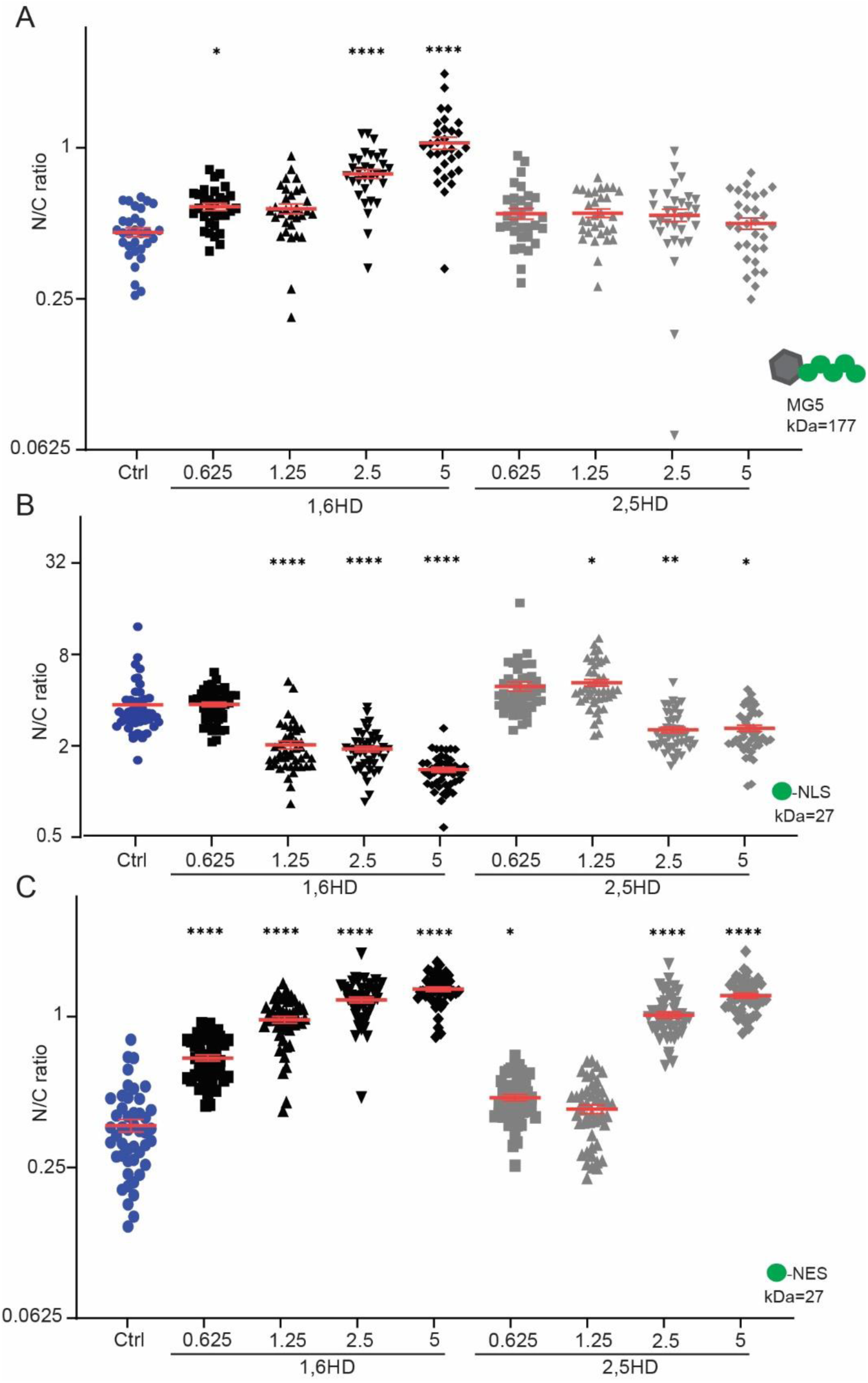
Disruption of NPC permeability barrier by 1,6HD. (A-C) Nuclear compartmentalization of GFP-based reporter proteins (MG5, GFP-NES, GFP-NLS) in yeast cells exposed for 10 min with the indicated concentrations of 1,6HD or 2,5HD. MG5 is a fusion of Maltose Binding Protein and 5 GFPs; GFP-NLS features the classical Simian Virus 40 NLS and GFP-NES the Stress-Seventy subfamily B1 NES. The N/C ratio is the ratio of the average fluorescence in the nucleus (N) over that in the cytoplasm (C). One-way ANOVA with Dunnett’s multiple comparison test comparing treatment to control was used to calculate the statistical significance of (A) MG5 and (C) GFP-NES, while the non-parametrical Kruskal-Wallis with Dunn’s multiple comparison test comparing treatment to control was used to calculate the statistical significance of (B) GFP-NLS. Error bars reflect SEM from the mean of three independent experiments. At least 30 cells per condition were analysed. P-values*<0,05 **<0,01 ****<0,0001.

To assess active import and export, GFP with a classical NLS (GFP-cNLS) and GFP-NES reporters are used. The balance between Kap60/Kap95-facilitated import of GFP-cNLS and its passive efflux leads to nuclear accumulation. Similarly, the balance of CRM1-facilitated export of GFP-NES and its passive influx leads to a steady-state nuclear exclusion. The import and export reporters showed a gradual decline in nuclear accumulation and exclusion, respectively, with increasing 1,6HD concentrations (Fig 1B,C). This loss of nuclear compartmentalisation could solely be the consequence of the increased passive permeability (Fig 1A), but could additionally be the result of a decrease in the active transport rates. As for passive transport 1,6HD had a stronger effect for active transport than 2,5HD, as higher concentrations of 2,5HD were needed to decrease the compartmentalisation of GFP-NLS and GFP-NES (Fig 1B,C). From this we conclude that exposure of live yeast cells to 1,6HD (10 min, 0-5%) leads to a gradual loss of the permeability barrier of NPCs.

### On the specificity of 1,6HD towards disrupting nuclear transport

The question if the increased NPC permeability after exposure to 1,6HD is a direct consequence of an altered nuclear transport system, or rather a consequence of indirect effects on the cell’s physiology, is pertinent. Indeed, depending on the exposure time and concentration 1,6HD may well have pleotropic effects in cells, as also previously discussed (Kroschwald, Maharana, and Simon 2017). Using the set concentration of 5% 1,6HD, we assessed all aspects of cell physiology that we deemed relevant and could assess. First, we treat the cells for 10 or 30 min with 5% 1,6HD or 2,5HD and observed no effects on cell viability (Fig 2A). Then, we assessed if 10 min exposure to 5% 1,6HD leads to changes in free ATP levels or cytosolic pH, using fluorescence-based sensors (Imamura et al. 2009; Miesenböck, De Angelis, and Rothman 1998). Our rationale for testing these was that ATP and pH levels could change when cells are experiencing metabolic stresses. We find, however, that the levels of free ATP are unchanged after 1,6HD treatment. As a control, sodium azide (NaN_3_) and 2-deoxy-glucose (2DG) were used, which both depleted the cell of energy (Fig 2B). The cytosolic pH values, calibrated as described in (Mouton et al. 2020), decrease mildly from 7.2 to 6,8 or 6,7 after exposure to 1,6HD and 2,5HD respectively, and therefore remain in the physiological range (Fig 2C).

**Figure 2:**
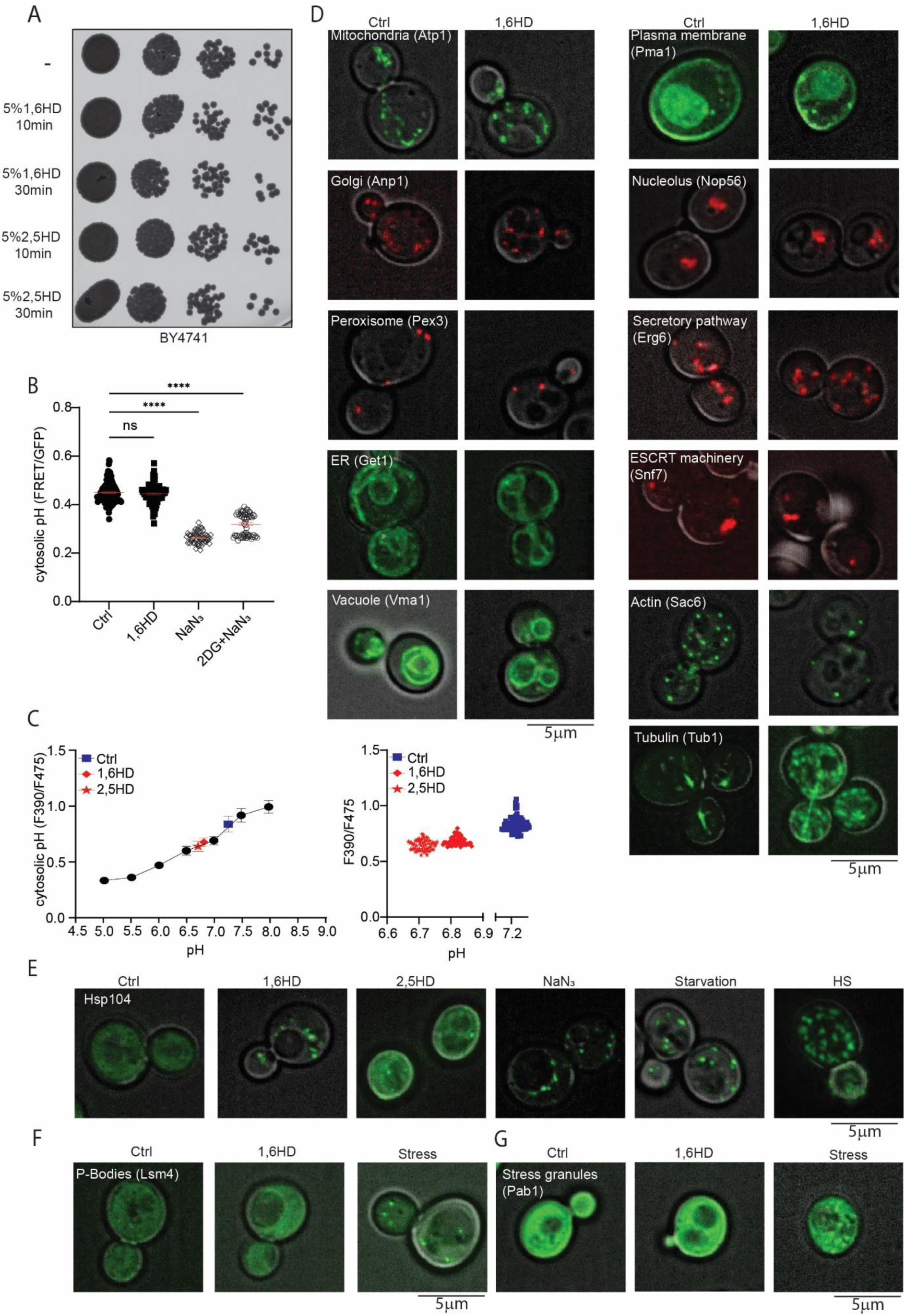
Impact of 1,6HD on cell survival, physiology and subcellular structures. (A) Growth assay showing serial dilutions of cultures exposed to 5% 1,6HD or 2,5HD for the indicated times. (B) Free ATP levels in cells measured using a FRET-based ATP-sensor; lower FRET/GFP ratio indicates lower free ATP. Cells were untreated (ctrl), exposed to 5% 1,6HD for 10 min, or exposed for 30 min to metabolic poisons azide (NaN_3_) or to NaN_3_ plus deoxyglucose (NaN_3_ + 2DG). The error bar of the scatter plot reflects SEM from the mean of three independent experiments. At least 60 cells per condition were analysed. Non parametric Mann-Whitney test was used to calculate statistical significance in FRET/GFP ratios comparing treatment to control. (C) Calibration curve for cytosolic pH values of the pH sensor pHluorin (F390/F475) in cells (black circles). The pH before (ctrl, blue squares) and after 10 min exposure to 1,6HD (red diamonds) or 2,5 HD (red stars) are indicated. Each point represents data from 60 cells (left grapgh), individual measurements are shown (right graph). (D) Fluorescence images of different cellular structures endogenously tagged with either GFP or mCherry, before and after 10 min exposure to 5% 1,6HD. (E) Fluorescence images showing localization of endogenously tagged Hsp104-GFP after 10 min exposure to 5% 1,6HD or 5% 2,5HD and under indicated stress conditions. (F,G) Fluorescence images showing localization of endogenously tagged Lsm4 (P-bodies, F) or Pab1 (Stress granules, G) with GFP after 10 min exposure with 5% 1,6HD and after induction of stress. Representative images of three independent replicates. The scales bars are 5μm.

Next, we looked at the morphology and localization of different subcellular structures using GFP- or RFP-tagged proteins marking the mitochondria, Golgi, peroxisome, ER, vacuole, plasma membrane, nucleolus, secretory pathway, and ESCRT machinery. From visual inspection we conclude there are no obvious changes in their appearance after 10 min exposure to 5% 1,6HD (Fig 2D). In contrast, the appearance of microtubules and actin filaments does change after treatment with 1,6HD, which aligns with some previous literature (Wheeler et al. 2016; Kroschwald, Maharana, and Simon 2017). Hsp104, a disaggregase that can refold and reactivate previously aggregated proteins and responds to alcohol-stress (Bösl, Grimminger, and Walter 2006; Sanchez and Lindquist 1990; Glover and Lindquist 1998; Harari et al. 2022), forms foci upon exposure to 1,6HD, similar to when cells are exposed to either nitrogen starvation, energy depletion or heat shock (Fig 2E), suggesting that 1,6HD induces some level of protein stress. Finally, 1,6HD does not induce the formation of p-bodies (Fig 2F) or stress granules (Fig 2G).

Taking the above together, under the conditions where mid exponentially growing cells are exposed to 5% 1,6HD for 10 min, there are effects on the cytoskeleton and Hsp104 to be noted, but cell viability, the pH and ATP levels in the cytosol, and the appearance of mitochondria, Golgi, peroxisomes, ER, vacuoles, plasma membrane, nucleolus, the secretory and ESCRT pathways and stress granules are not notably changed. While this is not an absolute proof of absence of indirect effects on nuclear transport, the data strongly suggest that the 1,6HD-dependent effects on NPC permeability shown in Fig 1 is due to direct effects on the nuclear transport machinery. Exposure to 10 min 5% 1,6HD thus permeabilizes NPCs with surprising specificity.

### 1,6HD induced loss of NTRs from the NPCs disrupts the permeability barrier

Previous work proposed that the effects of 1,6HD are related to the alcohol-sensitive hydrophobic interactions between the FG-nups that maintain the permeability barrier (Patel et al. 2007; Ribbeck and Görlich 2002; Schmidt and Görlich 2015). Indeed, when the FG-domains of Nup100 (Nup100FG) in preformed condensates are exposed to the concentrations of 1,6HD that were also used in life cells (0-5%), partial solubilisation of the condensates is observed (Sup fig 1). While disruption of FG-nup interactions by 1,6HD is indeed a scenario that is supported by *in-vitro* data, it is also one that is not easy to proof or disproof in *in vivo* experiments. Alternative or additional explanations for the increased permeability of NPCs in 1,6HD treated cells that can be experimentally addressed, relate to the composition of the NPCs and to the NTRs. We explore them both.

Previous work (Shulga and Goldfarb 2003) showed that 1,6HD did not lead to release of NPC components in wild type W303 cells, but it did in a mutant lacking Nup170. We noticed that even in wild type cells the exposure to 10% 1,6HD lead to release of NPC components (data not shown). Therefore, we repeated the analysis of nup localisation, and expanded on it with an analysis of proteins levels. We assessed the effects of 5% 1,6HD on the protein levels and NPC-association of nine representative endogenously tagged nups. The five tested FG-nups (Nsp1, Nup49, Nup159, Nup100, Nup116), two of the scaffold nups (Nup133 and Nup170) and two basket nups (Nup60 and Nup2) did not show changes in expression levels by western blot (Fig 3B). Also, their localization to the nuclear envelope was unchanged, consistent with (Shulga and Goldfarb 2003) (Fig 3B). We conclude that the 10 minutes 1,6HD treatment did not lead to dissociation or degradation of the tested NPC components, and hence it is unlikely that the increased permeability is a result of changes to the Nup-composition of the NPCs.

**Figure 3:**
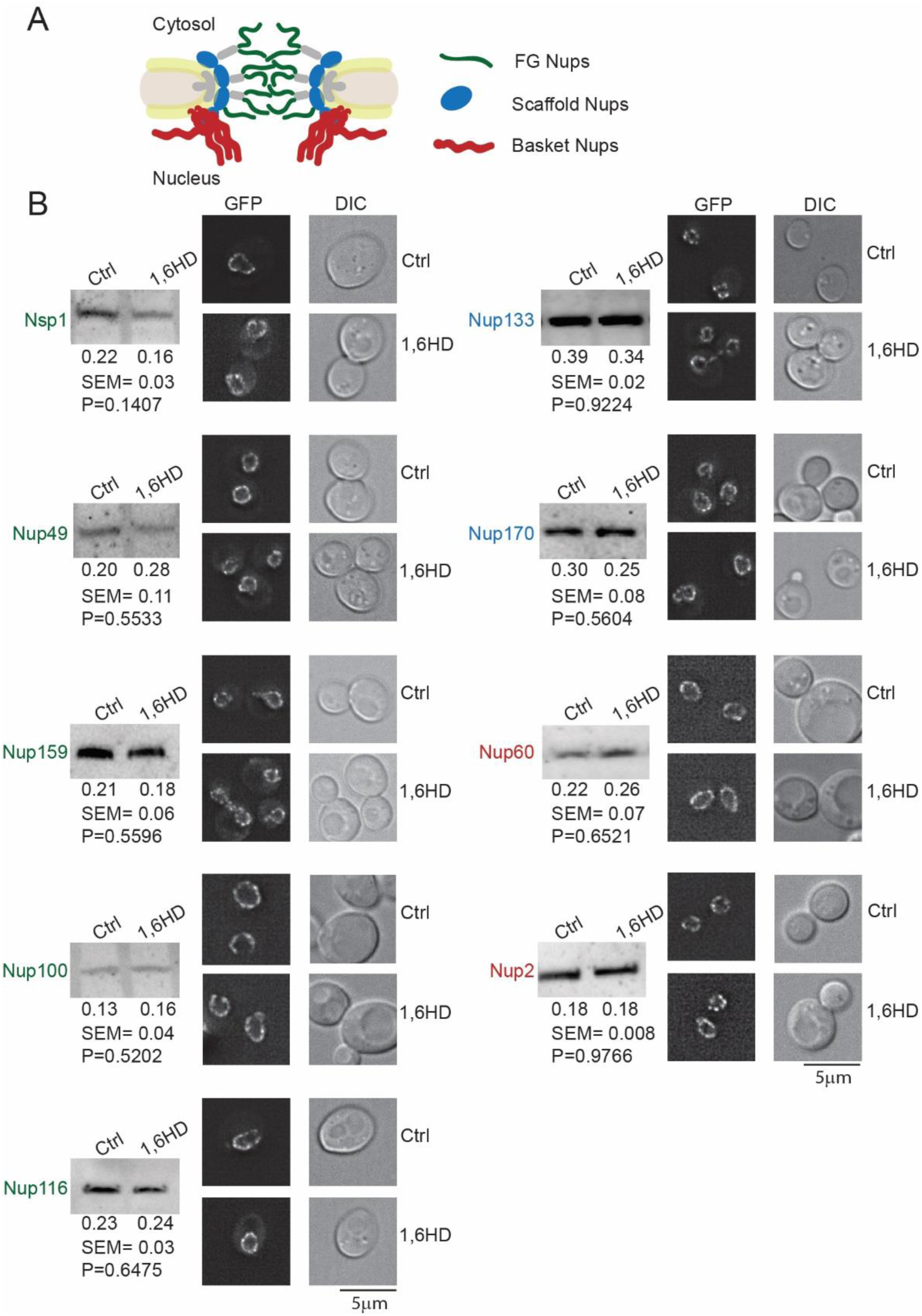
Impact of 1,6HD on the abundance and localization of NPC components. (A) Cartoon representation of NPC indicating the position of the nups analyzed in B. (B) Western blot of endogenous Nup-GFP protein levels before and after 10 min exposure to 5% 1,6HD; quantification gives mean, SEM and P values from at least three independent replicates. Fluorescence images of endogenously GFP-tagged nups after 10 min exposure with 5% 1,6HD. Representative images of three independent replicates. The scale bar is 5μm.

NPCs constitute a significant amount of NTRs at any point in time and their presence critically shapes the permeability barrier (Jovanovic-Talisman et al. 2009; Kalita, Kapinos, and Lim 2021; Lowe et al. 2015; Kim et al. 2018). Therefore, we addressed the localisation and abundance of endogenously GFP-tagged NTRs after treatment with 1,6HD. The interaction between the FG-nups and NTRs are based on dynamic multivalent binding with the phenylalanine’s of the FG-nups (Hoogenboom et al. 2021; Hough et al. 2015; Milles et al. 2015; Hayama et al. 2018; Sparks et al. 2018; Wing, Fung, and Chook 2022) and will thus also be sensitive to interventions disrupting hydrophobic interaction. We evaluated the localisation of endogenously GFP-tagged NTRs. Under normal conditions most NTRs are enriched at the nuclear envelope (NE) showing a punctate rim staining, e.g., Kap109, and few are enriched in the nucleus, e.g. Kap104 (Fig 4A). Strikingly, the exposure to 1,6HD led to a clear relocalisation of NTRs (Fig 4A). Kap104, Sxm1 (Kap108), Kap114, Nmd5 (Kap119), Pse1 (Kap121), Kap122 and Kap123 lose their accumulation at the NE or nucleus upon exposure to 1,6HD and distribute over the cytosol and nucleus (Fig 4A). Cse1 (Kap109), Kap120, Crm1 (Kap124) and Msn5 (Kap142) which are normally enriched at the NE, partly relocate. Kap60 and Kap95 were not visibly affected by the treatment probably related to the previously described immobile pool of Kap95 at NPCs (Lowe et al. 2015). Kap60 and Kap95 remain at NPCs while GFP-cNLS, whose active import is driven by Kap60-Kap95, loses nuclear accumulation (Fig 1B), suggesting that 1,6HD treatment increases passive permeability. When the less hydrophobic alcohol 2,5HD was used, it led to some NTRs losing their accumulation at the NE or nucleus, but always to a lesser extent compared to 1,6HD (Sup Fig 2). We conclude that the massive relocation of NTRs from NPCs may mechanistically explain the 1,6HD induced increase in the permeability of NPCs.

**Figure 4:**
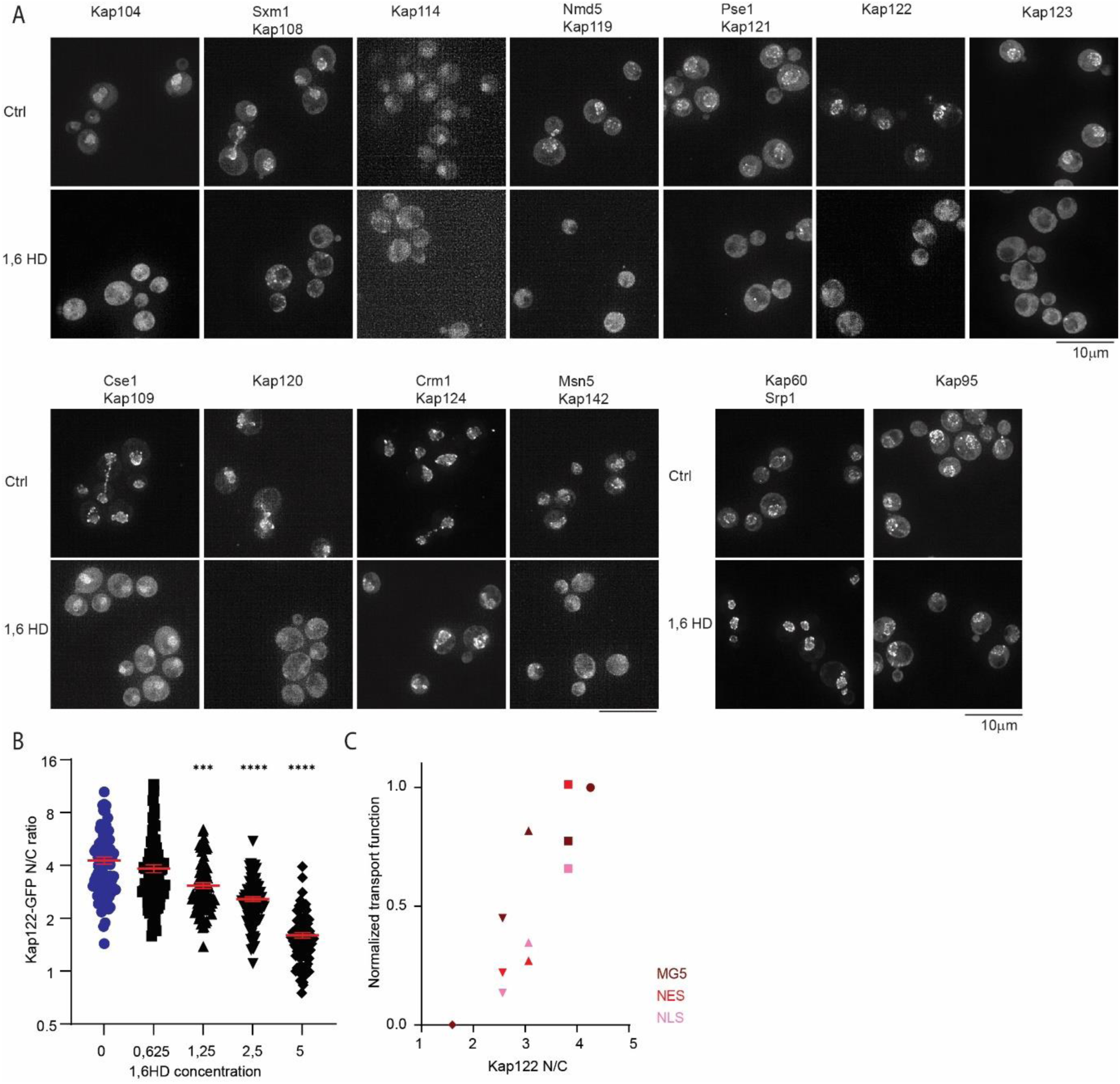
Impact of 1,6HD on NTRs. (A) Fluorescence images of endogenously GFP-tagged NTRs after 10 min exposure with 5% 1,6HD. Representative images of three independent replicates. The scale bar is 10μm. (B) Nuclear accumulation of Kap122-GFP in yeast cells exposed for 10 min with the indicated concentrations of 1,6HD. Non-parametrical Kruskal-Wallis with Dunn’s multiple comparison test was used to calculate statistical significance, comparing treatment to control. Error bars reflect SEM from the mean of three independent experiments. 90 cells per condition were analyzed. P-values ***<0,0005 ****<0,0001. (C) Average transport function measured with MG5 (dark red, normalized N/C from Fig1A), GFP-NLS (pink, normalized N/C from Fig1B) and GFP-NES (red, normalized N/C from Fig 1C) as a function of Kap122-GFP location at the NE and nucleus (from Fig 4B) under control conditions and increasing concentrations of 1,6HD (symbols as in 4B: 0% circles; 0,625% squares; 1,25% triangles up; 2,5% triangles down; 5% diamonds).

To further strengthen this interpretation, we sought to quantitatively correlate the concentration dependent NTR relocalisation, with the 1,6HD concentration dependent entry of the reporters used before: MG5 (Fig 1A), GFP-NLS (Fig 1B) and GFP-NES (Fig 1C). We chose Kap122 for this analysis as Kap122 clearly loses its accumulation at the NE and distributes over the cytosol and nucleus (Fig 4A). The localisation of endogenously tagged Kap122-GFP in the nucleus and NE was assessed in a strain co-expressing endogenously tagged Nup133-mCherry to mark the NE. The average nuclear accumulation of Kap122 gradually decreased from 4,3 to 3,8 to 3,1 to 2,6 to 1,6 upon exposure to zero, 0.625, 1.25, 2.5 or 5% 1,6HD. Moreover, we could correlate Kap122 relocalisation from the nuclear envelope (NE) under these conditions with the measured passive permeability of NPCs for MG5 (Fig 4C), GFP-NLS (Fig 4C) and GFP-NES (Fig 4D) with a Pearson correlation coefficient of 0.9, 0.8 and 0.9 respectively. These correlations support that 1,6HD perturbs the NPC permeability barrier by releasing the NTRs.

## DISCUSSION

Here we assessed the specificity and mechanism by which 1,6-hexanediol (1,6HD), an aliphatic alcohol that interferes with hydrophobic interactions, disrupts the permeability barrier of NPCs in live baker’s yeast cells. Exposure of live yeast cells to 1,6HD (10 min, 0-5%) leads to a gradual loss of the permeability barrier of NPCs. We conclude this is likely a direct effect on the nuclear transport machinery as cell viability, the pH and ATP levels in the cytosol, and the appearance of mitochondria, Golgi, peroxisomes, ER, vacuoles, plasma membrane, nucleolus, secretory pathway and stress granules were not notably changed. There were effects on the cytoskeleton and protein homeostasis (Hsp104 foci) to be noted and we cannot exclude that 1,6 HD impacts the cell’s physiology in ways that we did not monitor. Mechanistically we propose that the displacement of NTRs from the NPC underlies the loss of NPC function because 1,6HD treatment induced a massive relocation of multiple NTRs from NPCs. This displacement from the nuclear envelope quantitatively correlated with the passive permeability of NPCs.

Our studies align well with previous reports that showed that the selective properties of the FG-nups rely on the physical presence of NTRs within the NPC. The earliest study is one showing that the presence of transport factor enhances the selectivity of FG-nucleoporin-coated membranes (Jovanovic-Talisman et al. 2009). The most recent reports on detergent-permeabilized human cells show that the enrichment of NTRs at the NPCs is important for the permeability barrier by preventing passive permeability (Kalita et al. 2022). Our work adds to this by showing the importance of NTRs in live cells. The benefit being that in live cells there is a constant and large flux of transport and therefore, together with the loss of the estimated 15,6 MDa of NTRs from the central channel also 10,4 MDa worth of cargo is being lost (Kim et al. 2018). This joint loss of NTRs *and* cargo from the NPC central channel will present a major change in the macromolecular crowding and composition, and hence its physicochemical properties. How this alters the structural dynamics of the FG-nups, and if this poses a risk for NPC function would be interesting questions for the future.

Extrapolating from studies using purified FG-nup fragments that proposed that the effects of 1,6HD is related to the alcohol-sensitive hydrophobic interactions between the FG-nups (Patel et al. 2007; Ribbeck and Görlich 2002; Schmidt and Görlich 2015) one may expect that 1,6HD also alters the interactions between the FG-nups in our assays using live cells. This is, however, difficult to address in live cells. Hence it remains unclear if the NTRs are released from the NPCs as a consequence of a lowered binding affinity between FG-nups, or because 1,6HD directly lowered the binding affinity of NTRs for the FG-repeat regions. If one considers that the functional composition of central channel is a system composed of NTRs *and* FG-nups in close collaboration, then the discrimination between these scenarios becomes less important.

An unanswered question in the field is if NPCs that are dysfunctional can be detected and removed. To assess this question, one needs to be able to inducibly damage NPCs. NPC permeabilization is expected to be an intervention that triggers quality control similar to when assembly fails (Thaller et al. 2019; Webster et al. 2016; Thaller et al. 2021). The here described method could provide a tool to study the recruitment of quality control factors and to follow the repair or degradation.

Lastly, our study may serve as a warning that the effects of 1,6HD on liquid-liquid phase separation of diverse cellular macromolecular complexes may actually be the consequence of to 1,6HD’s prime effect on the NPC and cognate NTRs. We speculate that the hydrophobic and highly acidic nature of NTRs may readily compromise their stability above a critical concentration. Consistent with this is that overexpression of Sxm1, Kap95, and Kap114 is toxic to cells (Semmelink et al. 2022). In any case, a major misplacement of NTRs and associated cargo will dramatically change the nuclear and cytoplasmic proteomes and this may generally compromise their stability. The increase in the number of Hsp104 foci that we observe may indeed reflect such loss of protein homeostasis.

Altogether, this paper puts hydrophobic interactions between NTRs and FG-Nups centre stage in the explanation of the selective properties of NPCs supporting the Kap-centric model for nuclear transport proposed by the Lim laboratory (Springhower, Rosen, and Chook 2020).

**Table 1:**
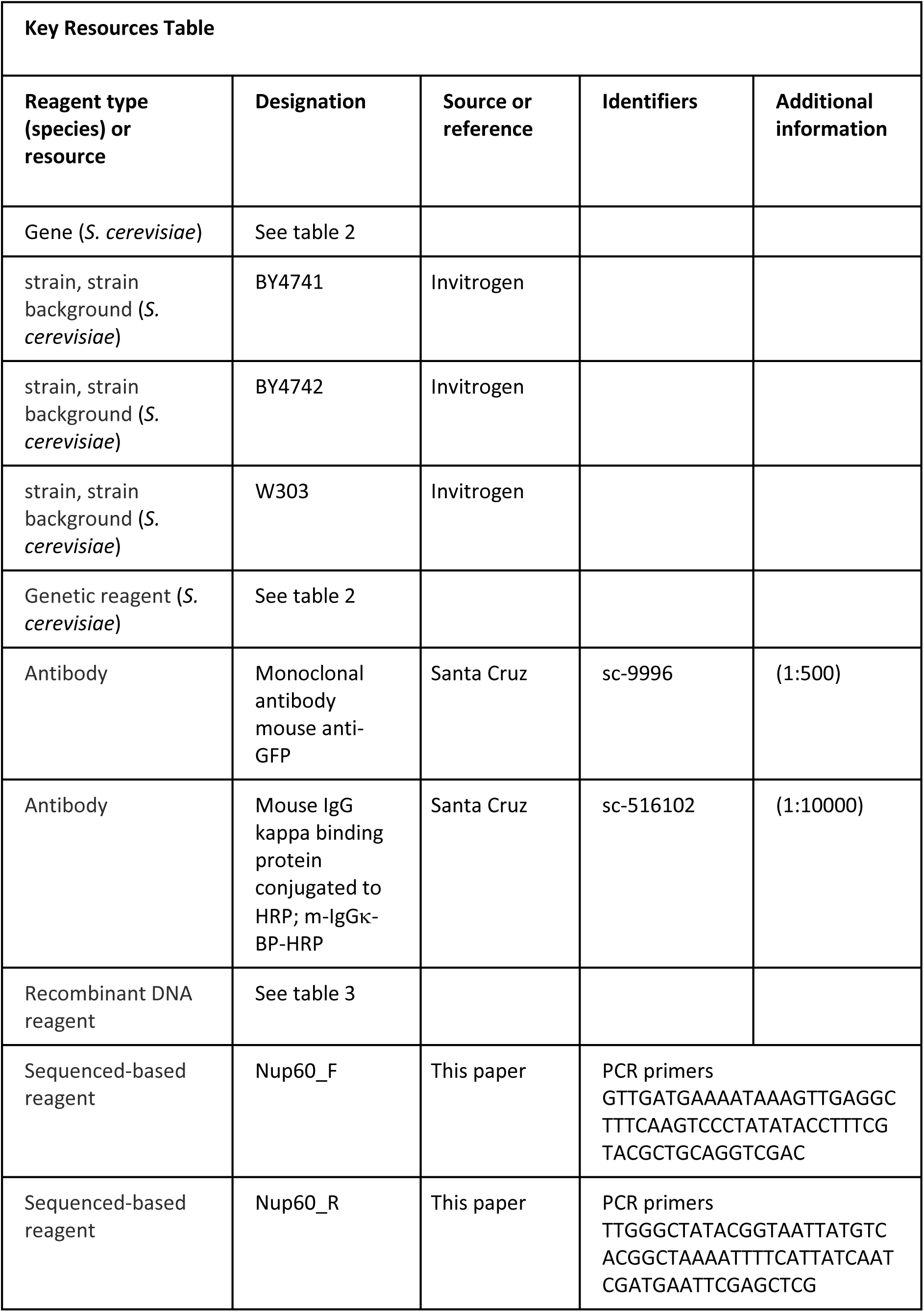
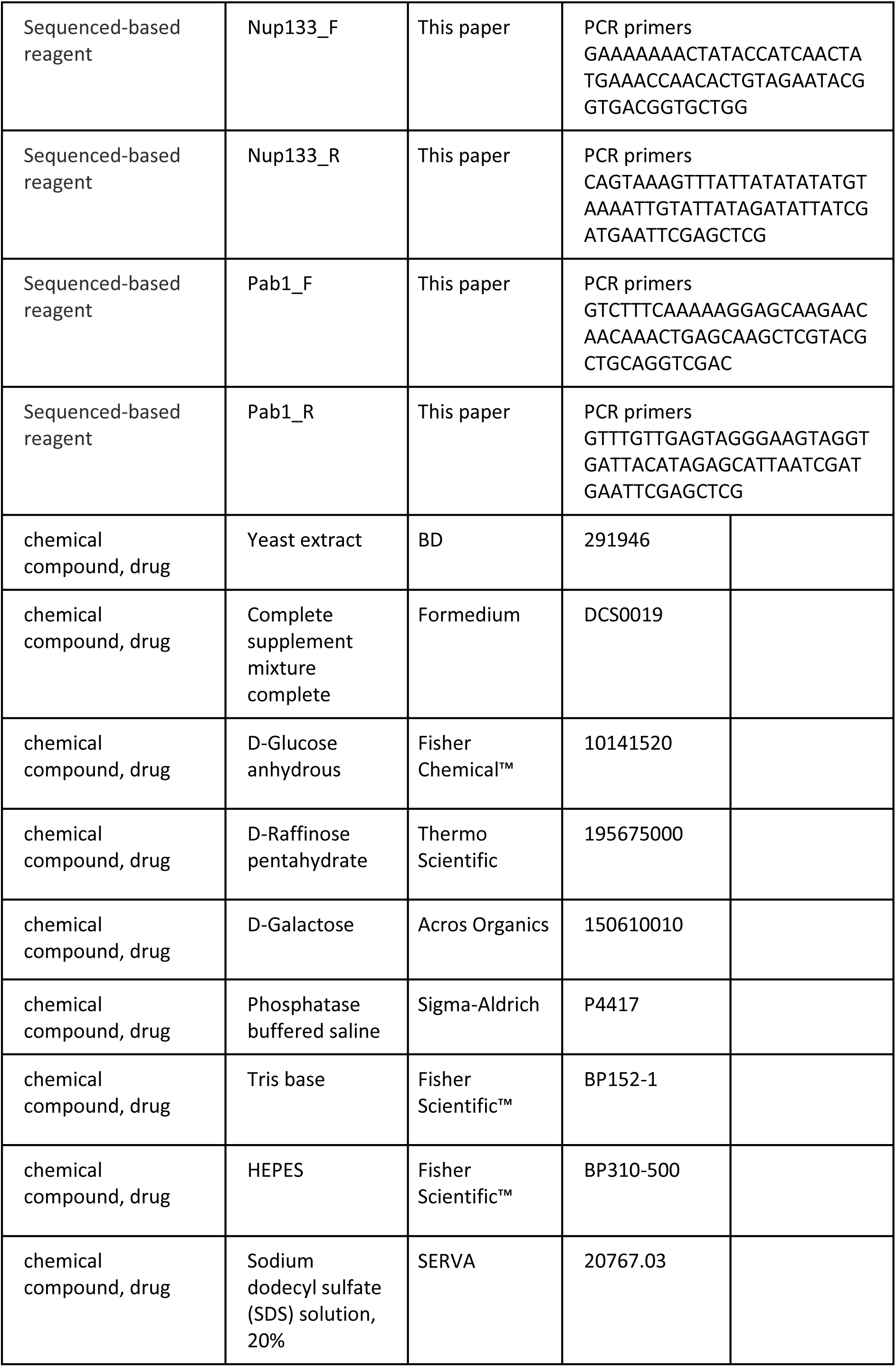
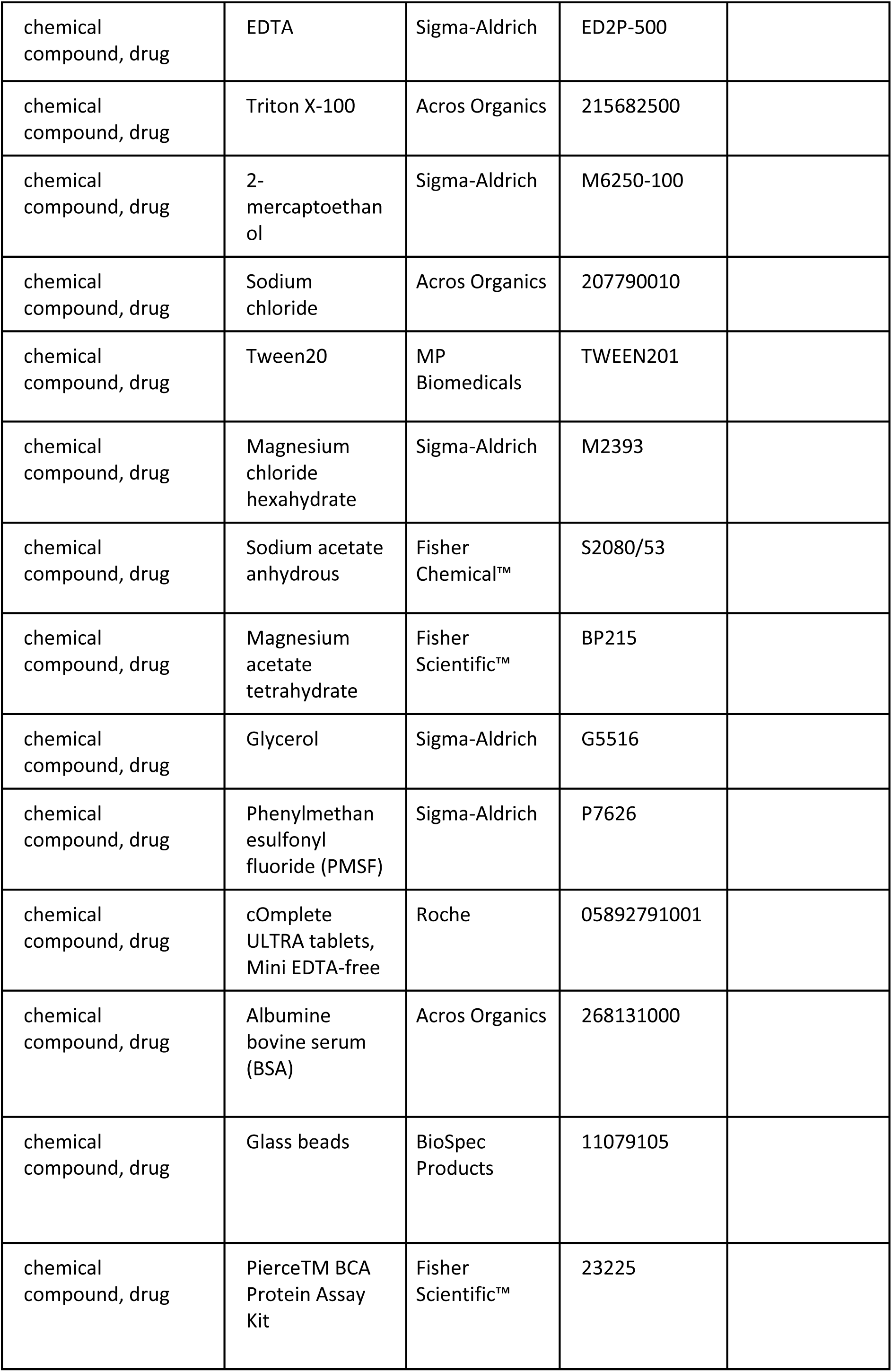
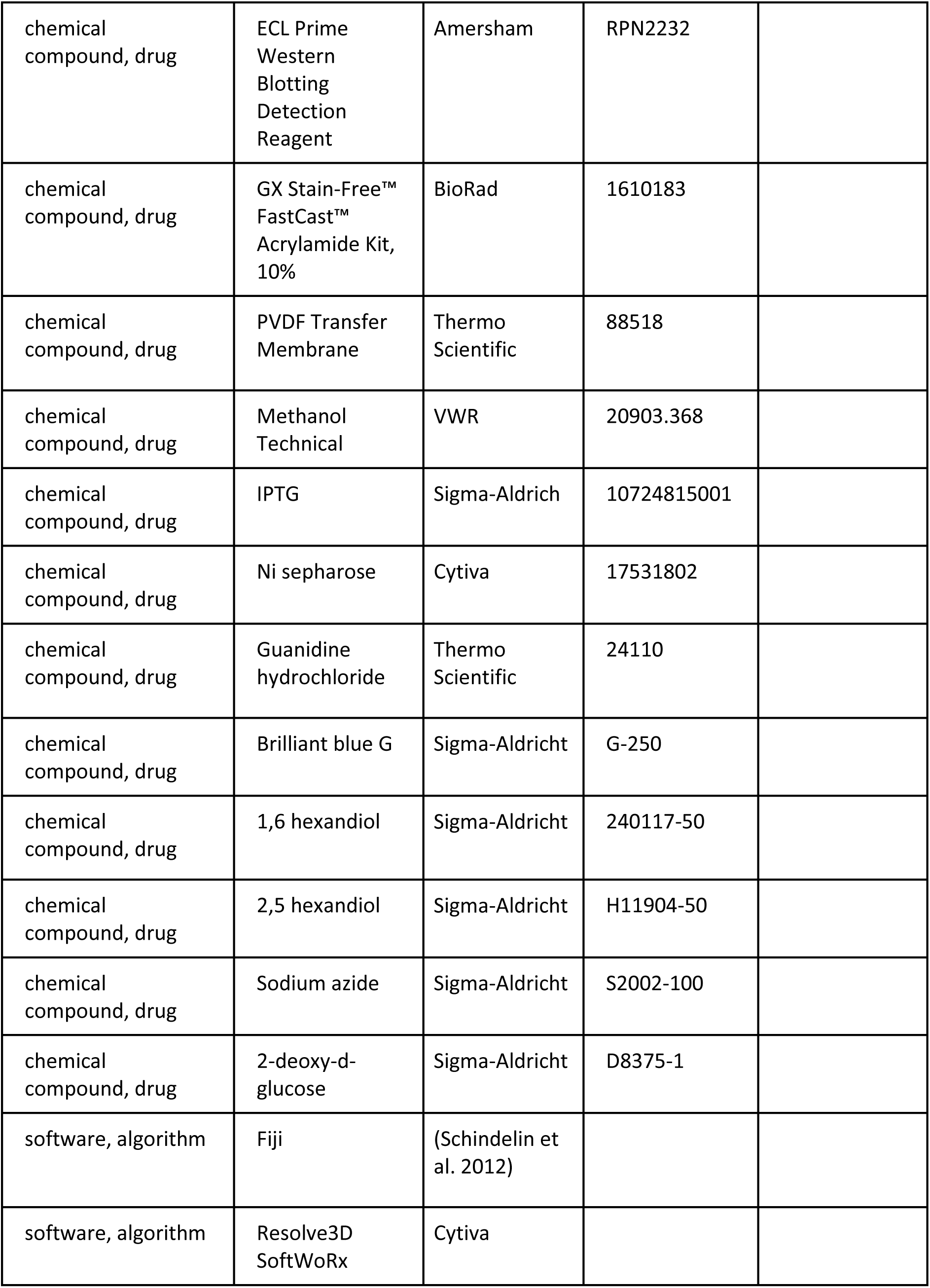
Key resources table

## MATERIAL AND METHODS

### Strains and Growth conditions

All *Saccharomyces cerevisiae* strains used in this study have the BY4741 background, except yER016, which were created in the W303 background. Strains are listed in Table 2. yER016, yER020 and yER023 were created as described in (Janke et al. 2004). GFP-tagged strains were taken from the 4000-GFP yeast library (Thermofisher), RFP-tagged strains were taken from the localization database collection (Huh et al. 2003).

**Table 2.**
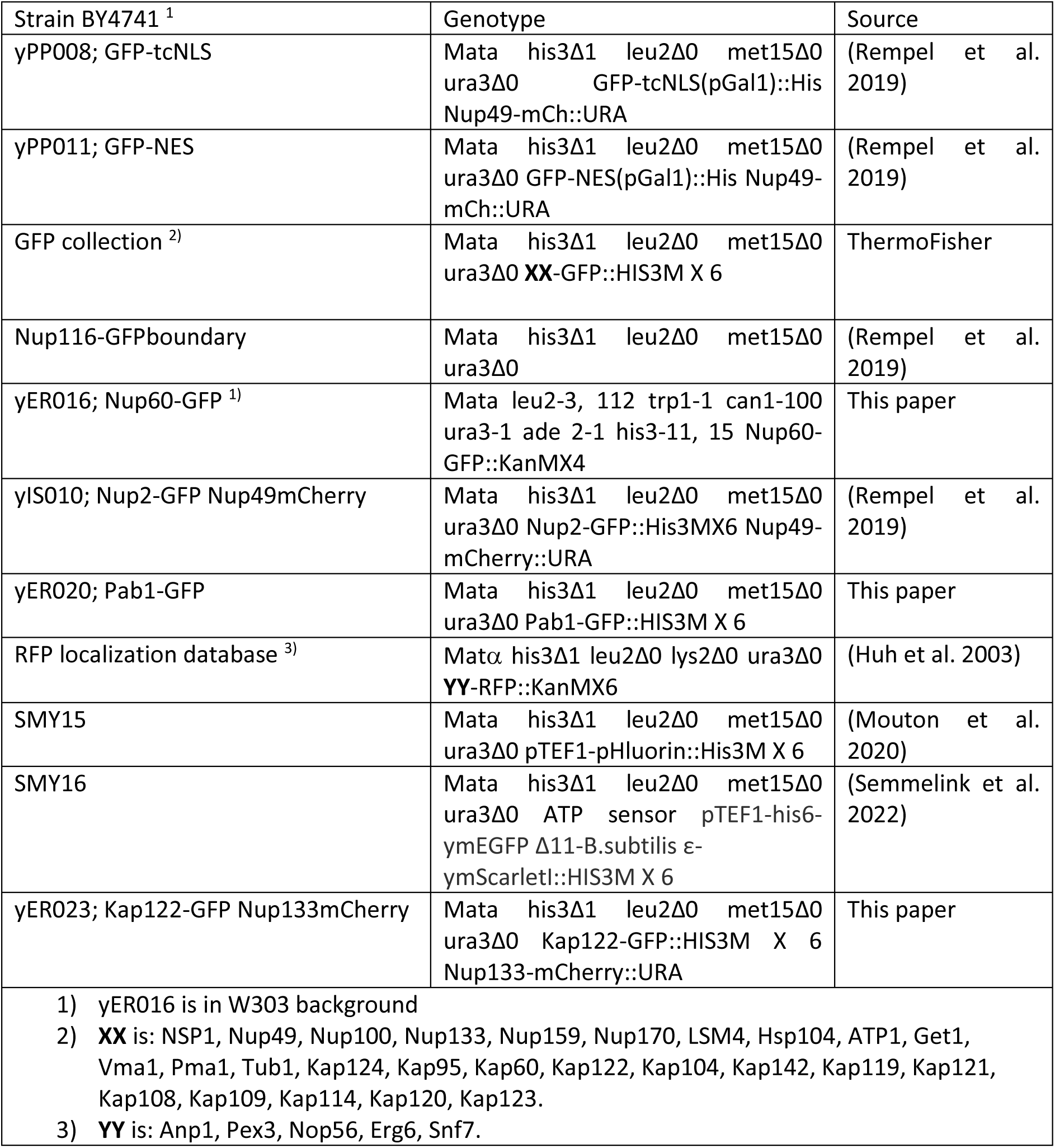
Yeast strains used in this publication

Cells were grown at 30°C, with shaking at 200 RPM on Synthetic Complete (SD) medium supplemented with 2% (w/v) glucose. Cells from an overnight culture were diluted 1:10 during the day and then again for an overnight culture in SD-2% glucose. Cells were diluted again on the day of the experiment, and grown for several hours to obtain cultures in exponential growth phase (OD_600_ 0.6-0.8) before each experiment.

### Spot assay

On the day of the experiment, exponentially growing cells were treated with 5% 1,6HD or 5% 2,5HD for 10 or 30 minutes, as indicated in Fig. 2A, and diluted in sterilized milliQ water to obtain 10^6^ cells/ml, and further serial diluted in milliQ water. 5μl of each dilution was spotted on YPD plates and the plates were imaged after 48H growth at 30°C.

### Microscopy

All *in vivo* experiments were performed at 30°C. Images were acquired using a DeltaVision Elite imaging system (Cytiva) composed of an inverted microscope (IX-71; Olympus) equipped with a UPlanSApo 100x (1.4 NA) oil immersion objective, InsightSSI solid-state illumination, and an EDGE sCMOS 5.5 camera. For all experiments, stacks of 30 images with 0.2μm spacing were taken.

### Protein lysate and Western Blot

20 ml of yeast culture was grown to an OD_600_ 0.8-1.2. Cells were subsequently treated with 5% 1,6HD for 10 min at 30°C, with shaking at 200 RPM. After the treatment, cells were centrifuged, and all the following steps were performed at 4°C. The cell lysate was resuspended in 0.25ml of lysis buffer (50mM HEPES, 200mM sodium acetate, 1mM EDTA, 5mM magnesium acetate, 5% glycerol, 1% triton x-100, 10mM β-mercaptoethanol, protease inhibitor without EDTA) and lysed in two rounds of bead-beating in a Fastprep device (MP biomedicals). Lysates were cleared by consecutive centrifugations at 6000 x g for 5 min, followed by centrifugation of the supernatant at 17700 x g for 5 min. The resulting supernatant was centrifuged once more at 17700 x g.

Western blots were performed as follows: whole cell lysates were separated by SDS-PAGE. The proteins were subsequently transferred to PVDF membranes. After blocking with 5% skim milk in TBS-T, GFP-tagged proteins were detected with anti-GFP (Santa Cruz sc-9996 HRP) was used, followed by HRP-conjugated mouse IgG kappa binding protein (Santa Cruz sc-516102, m-igGκ BP-HRP).

### Expression and purification of nucleoporin FG-domains

Nup100FG domains were expressed and purified as described in (Kuiper et al. 2022). In short: FG-domains proteins with an N terminal His-tag and a unique C-terminal cysteine were expressed in *Escherichia coli*, by induction with 0.5mM IPTG and purified from cell extracts on a Nickel-Sepharose column under denaturing conditions (2M GuHCl, 100mM Tris-HCl pH 8). The C-terminal cysteine was reduced with DTT and blocked by modification with Iodoacetamide. Protein purity was checked with SDS-PAGE and subsequent Brilliant Blue staining.

### Spin Assay

A concentrated stock of 100μM Nup100FG domains in 2M GuHCl, 100mM Tris-HCl pH 8, was diluted to 3μM into TBS (50mM Tris-HCl, 150mM NaCl pH 8). The protein was left to self-assemble into particles for 1h at RT, and then the protein was treated for 10 min with different concentrations of 1,6HD. Samples were centrifuged (17.700 x g for 10 min at RT), and soluble and insoluble fractions were run separately on SDS PAA gels. Gels were stained with Brilliant Blue G (Sigma-Aldrich, G-250) and imaged using a BioRad chemidoc (BioRad). Band intensities were determined using Fiji (Image J, National Institute of Health).

### Determining the intracellular pH with the pHluorin sensor

pHluorin ratios were calibrated in live cells in buffers with a pH of 5, 5.5, 6, 6.5, 7, 7.5, and 8, as described in (Mouton et al. 2020). The FRET/CFP and FRET/mEGFP (F390/F475) ratios were determined from cells on a glass slide. Cells were then treated with 1,6HD as described in Fig 2, and a calibration curve was used to determine the pH change after treatment.

**Table 3.** Plasmids used in this publication.

| Plasmid number | Genotype | Source |
| --- | --- | --- |
| pPP008; MG5 | pUG34-Gal1-MBP-5XGFP-His | (Popken et al. 2015) |
| pACM063; mCh-L-TM | pUG36-Gal-mCherry linker-TM-URA | (Meinema, Poolman, and Veenhoff 2013) |
| pYM28 | pAgTEF-SpHIS5-tAgTEF | Euroscarf, Janke et al 2004 |
| pYM30 | pAgTEF-kanMX-tAgTEF | Euroscarf, Janke et al 2004 |
| pPP014 | mCherry-Ura cassette | (Rempel et al. 2019) |

### ATP sensor values and free ATP levels

Cells expressing a FRET-based ATP sensor (Semmelink et al. 2022), were used to determine free ATP levels as described in (Semmelink et al. 2022). Cells were treated as described in Fig 2, imaged, and the FRET over GFP ratio was calculated using Fiji (see below).

### Image Analysis

All images were processed using Fiji (Image J, National Institute of Health). For each image, the z-stack with the best focus was selected. For GFP-tagged reporters, we determined the fluorescence around the nuclear envelope and subtracted the background from outside the cell. For pHluorin and the ATP sensor, we determined the fluorescence in each channel for each cell and took the fluorescence of the entire cell and subtracted the background from a region outside the cell for each channel. The respective ratios were subsequently calculated. To quantify the nuclear localization (N/C ratio) of the GFP-based reporters and Kap122, the average fluorescent intensity of the nucleus and the cytosol was measured. The nucleus area was determined using either the mCherry-TM reporter (pACM063) that indicated the nuclear envelope (Fig 1) or Nup133-mCherry (Fig 4B). A section of the cytosol excluding the vacuole was selected to measure the fluorescence in the cytosol.

### Statistical Analysis

Statistical parameters, including the number of cells analyzed, are reported in figure legends. All regressions and correlations leading to the sigmoidal curve equation, R^2^, and all Pearson’s correlation statistics were done in GraphPad Prism.

## ACKNOWLEDGEMENTS

ERB and TO are supported by PhD-fellowships from the Graduate School of Medical Sciences of the University of Groningen. ERB, AS, LMV, are supported by a Vici grant (VI.C.192.031) from the Netherlands Organisation for Scientific Research. We want to thank Amarins Blaauwbroek for practical assistance.

## AUTHOR CONTRIBUTIONS

ERB and LMV conceived the project. ERB designed, performed and analysed all experiments with help from SNM (Fig. 2BC) and TO (Supfig1). The manuscript was written by ERB and LMV with input of all authors.

## DATA AND REAGENT AVAILABILITY

All data and reagents are available upon request.

---

**Supplementary Figure 1.**
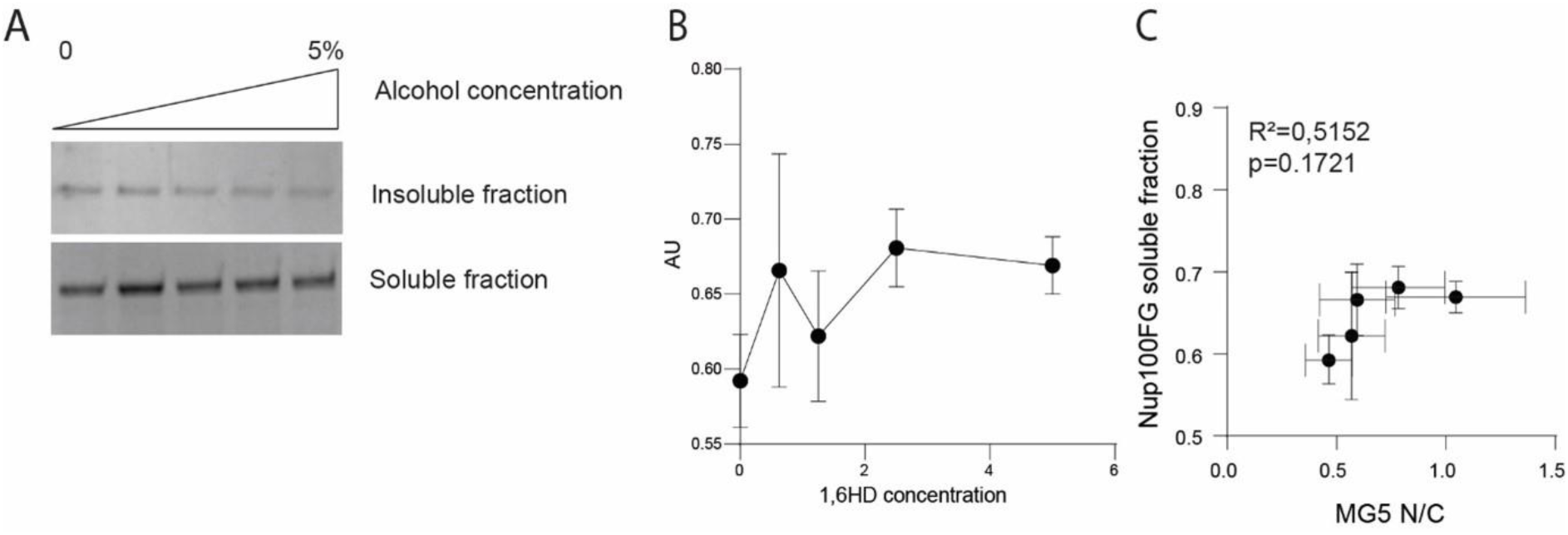
(A) Purified Nup100FG domains were left to form condensates for 1 hour and subsequently treated for 10min with 0, 0.625, 1.25, 2.5 or 5% 1,6HD. Soluble and insoluble fractions were obtained by centrifugation, separated by SDS-PAGE and visualized by Brilliant Blue staining. Representative image of three independent experiments. (B) Quantification of the soluble fractions in (A) Error bars reflect SEM of three independent experiments. (C) Pearson correlation coefficient and two-tailed P values were calculated for the N/C ratio of reporter MG5 against the soluble fraction of Nup100FG domain after different concentrations of 1,6HD. Error bars reflect SEM from the mean of three independent experiments.

**Supplementary Figure 2.**
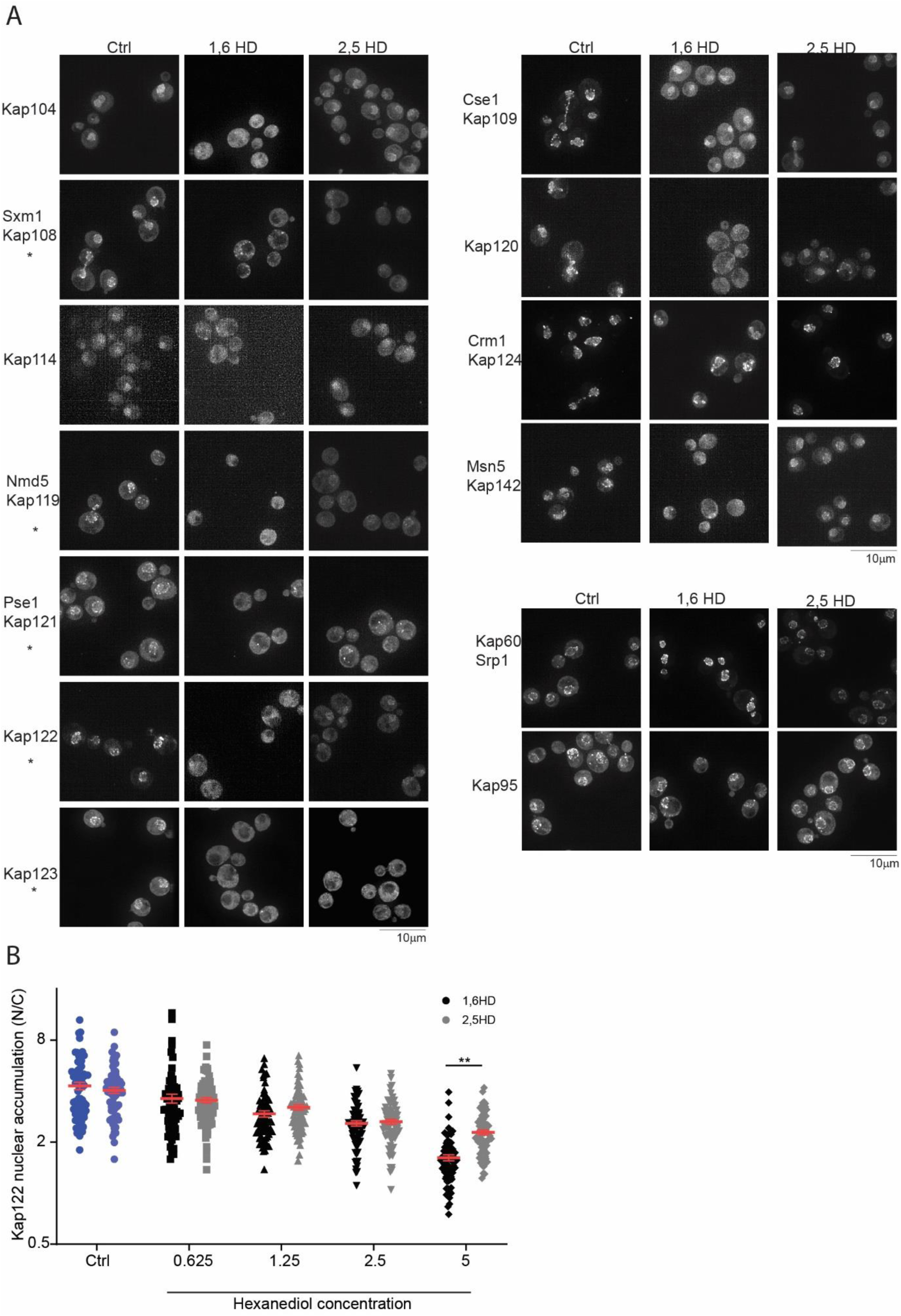
(A) Fluorescence images of endogenously GFP-tagged NTRs after 10 min exposure with either 5% 1,6HD (middle, as I Fig 4A) or 5% 2,5HD (right). Representative images of three independent replicates. The scale bar is 10μm. (B) Nuclear accumulation of Kap122-GFP in yeast cells exposed for 10 min to the indicated concentrations of either 1,6HD (as in Fig 4B) or 2,5HD. Non-parametrical Kruskal-Wallis with Dunn’s multiple comparison test comparing treatment to control was used to calculate statistical significance. Error bars reflect SEM from the mean of three independent experiments. 70 cells per condition were analyzed. P-values **<0,005.

